# Alleviatory effects of Danshen, Salvianolic acid A and Salvianolic acid B on PC12 neuronal cells and *Drosophila melanogaster* model of Alzheimer’s disease

**DOI:** 10.1101/2020.05.12.089797

**Authors:** Florence Hui Ping Tan, Andrew Chung Jie Ting, Nazalan Najimudin, Nobumoto Watanabe, Ghows Azzam

**Affiliations:** School of Biological Sciences, Universiti Sains Malaysia, 11800 Penang, Malaysia; USM-RIKEN International Centre for Ageing Science (URICAS), Universiti Sains Malaysia, 11800 Penang, Malaysia; Bioprobe Application Research Unit, RIKEN Centre for Sustainable Resource Science, RIKEN, Japan

**Keywords:** Danshen, Salvianolic acid A, Salvianolic acid B, *Drosophila melanogaster*, Alzheimer’s disease

## Abstract

Alzheimer’s disease (AD) is the most common form of neurodegenerative disorder worldwide. Its pathogenesis involves the hallmark aggregation of amyloid-beta (Aβ). Of all the Aβ oligomers formed in the brain, Aβ42 has been found to be the most toxic and aggressive. Despite this, the mechanism behind this disease remains elusive. With the ability to utilize various genetic manipulations, *Drosophila melanogaster* is ideal in analysing not only cellular characteristics, but also physiological and behavioural traits of human neurodegenerative diseases. Danshen water extract (DWE), obtained from the root of *Salvia miltiorrhiza* Bunge, was found to have a vast array of beneficial properties. In this study, DWE, and its major components, Salvianolic acid A (SalA) and Salvianolic acid B (SalB) were tested for their abilities to ameliorate Aβ42’s effects. DWE, SalA and SalB were confirmed to be able to reduce fibrillation of Aβ42. As Aβ42 causes neurodegeneration on neurons, DWE, SalA and SalB were tested on Aβ42-treated PC12 neuronal cells and were shown to increase cell viability. DWE and its components were then tested on the *Drosophila melanogaster* AD model and their rescue effects were further characterized. When human Aβ42 was expressed, the *Drosophila* exhibited degenerated eye structures known as the rough eye phenotype (REP), reduced lifespan and deteriorated locomotor ability. Administration of DWE, SalA and SalB partially reverted the REP, increased the age of AD *Drosophila* and improved most of the mobility of AD *Drosophila*. In conclusion, DWE and its components may have therapeutic potential for AD patients and possibly other forms of brain diseases.

## Introduction

The natural progression of aging has been a risk factor to age-related ailments such as dementia, with Alzheimer’s disease (AD) being the most common form (Ferri et al., 2005; Hung et al., 2010). AD is a neurodegenerative disease clinically depicted as a gradual and progressive decline in cognitive function (Murphy & LeVine, 2010). AD patients often encounter a range of symptoms such as behavioural changes to motor deterioration, and ultimately the inability to perform the simplest tasks. There are several hypotheses behind the occurrence of AD, with the two most established hypotheses being amyloid aggregation and tauopathy (Ittner & Götz, 2011; Tan & Azzam, 2017). Currently there are a few United States Food and Drug Administration (FDA)-approved drugs, however, all of which could only temporarily lessen AD symptoms. The inadequate understanding on the exact cause of AD affects the advancement of effective drugs (Knopman, 2006). Hence, are we looking into compounds that can potentially treat AD.

Here, we are using *Drosophila melanogaster* as the model organism to test the compounds of interest as it has been extensively utilized to study human disorders which includes neurodegenerative diseases such as AD (Moloney et al., 2010; Tan & Azzam, 2017). A key feature of using *D. melanogaster* is its short lifespan, requiring about 10 days to reach adulthood from egg (Helfand & Rogina, 2003; Sun et al., 2013; Tan & Azzam, 2017). The simple anatomy as well as genetic characteristics of *D. melanogaster* further supports its role as an ideal model for diseases. While it has fewer genes compared to *C. elegans, D. melanogaster* has 196 out of 287 recognised human disease genes homologues (St Johnston, 2002; Tan & Azzam, 2017). Furthermore, gene characterisation in *D. melanogaster* is simpler due to it having less genetic redundancy compared to vertebrate models. Despite having a much simpler structured brain, it possesses similar characteristics to the central nervous systems of mammals. *D. melanogaster* also exhibit age-dependant behaviours and many cellular processes which are involved in neurodegeneration. (McGurk et al., 2015; Tan & Azzam, 2017).

The dried root of red sage (*Salvia miltiorrhiza*) or generally known as Danshen is a popular traditional Chinese medicine used in clinical applications for over 1, 000 years (Chen et al., 2014). Its popularity is credited to its usage in the treatment of many diverse diseases such as AD, Parkinson’s disease, cerebrovascular disease as well as coronary heart disease (Zhou et al., 2005; Su et al., 2015). With over 100 isolated components (Pang et al., 2016), Danshen has a huge array of secondary metabolites with two dominant secondary metabolites classes; diterpenoids and phenolic acids (Su et al., 2015; Hügel & Jackson, 2014; Mei et al., 2019). Phenolic acids which include Salvianolic Acid A (SalA) and Salvianolic Acid B (SalB) to name a few, possess wide biological activities such as anti-oxidative, anti-coagulation as well as anti-inflammatory (Mei et al., 2019; Jiang et al., 2005). In this study, we will look at the effect of Danshen water extract, and its components SalA and SalB, in elucidating their protective roles against Aβ42-associated neurodegeneration using *Drosophila* AD model.

## Materials and Methods

### Compounds

Danshen water extract (DWE), Salvianolic acid A (SalA) (CAS no.: 96574-01-5) and Salvianolic acid B (SalB) (CAS no.: 115939-25-8) were obtained from LifeTech Solution Venture, Malaysia and were prepared in 0.5% dimethyl sulfoxide (DMSO) unless stated otherwise. Purities for SalA and SalB were over 98%. HPLC-grade formic acid was obtained from Merck while HPLC-grade acetonitrile was bought from Fisher Chemical.

### Preparation of Aβ42

Aβ42 was purchased from Anaspec (Cat no.: AS-20276) and 1 mg of the peptide dissolved in 1 mL of 100% DMSO. The peptide was kept in aliquots of 50 μL at −80 °C until further use.

### Thioflavin T (THT) Aβ42 Aggregation assay

ThT assays were performed using the SensoLyte ® Thioflavin T β-Amyloid (1 – 42) Aggregation Kit (Anaspec, Cat no: AS-72214) with slight modifications to the manufacturer’s protocol. In a black 96 well-plate, Aβ42 (Volume: 42.5 μl, final concentration = 42.5 μM) was added to the compounds (Volume: 2.5ul, final concentration = 1 mg/ml). Thioflavin T (Volume: 5 μl, final concentration = 20 μM) was the added in a dark room. The fluorescence reading was read using the Biotek Synergy 2 SLFP Multimode Microplate Reader every 300 seconds for a total of 3600 seconds at an excitation/emission of 440 nm/484 nm with pulsed shaking in between at 37 °C. The commercially available compound Morin (Volume: 2.5 μl, final concentration = 50 μg/ml) was used as the positive control.

### PC12 cell culture husbandry

Rat pheochromocytoma PC12 cells were obtained from the RIKEN Cell Bank (Tsukuba, Ibaraki, Japan). The cells were cultured routinely in Dulbecco’s Modified Eagle Media (DMEM) (Gibco, Cat no.: C11995500BT) supplemented with 10% Fetal bovine serum (Sigma Chemical Co. St Louis, MO, USA, Cat no.: 172012) and 10% Horse serum (Gibco, Cat no.: 26050-070) in a humidified atmosphere of 5% CO2 at 37 °C. To differentiate PC12 neuronal cells, cultures were provided with DMEM containing 10% Horse serum and 100 ng/mL Nerve Growth Factor (Sigma Chemical Co. St Louis, MO, USA, Cat no.: H9666-10UG).

### Cell viability determination by ATP assay

Aβ was added with respective compounds and aged for 72 hours at 37°C (Moreira et al., 2007). Cells were seeded in white 96 clear bottom well plates at 5×10^3^ densities per 100 μl with complete media. After 24 hours, the media was replaced with differentiation media and cells were incubated for 72 hours. Aβ (Final concentration = 10 μM) with respective compounds (Final concentration = 50 μM) were added to the wells and incubated for 24 hours. Equal amounts of CellTiter-Glo (Promega, Cat no: G7570) were added to the wells. The plate was shaken for 2 minutes and was left to stand in room temperature for 10 minutes. Luminescence reading was taken using Varioskan™ LUX multimode microplate reader. The experiment was done in triplicates.

### *Drosophila* Stocks and Husbandry

All *Drosophila* stocks used in this study are listed in Flybase (http://fybase.bio.indiana.edu). The following stocks were obtained from Bloomington *Drosophila* Stock Center (Bloomington, U.S.A.): Oregon-R wild type (#5), Glass Multiple Reporter-GAL4 (#1104) and UAS-Aβ42 (#33769) while Actin5C-GAL4 (#107727) was obtained from Kyoto Drosophila Genome and Genetic Resources (KGGR). All stocks were raised at 25 °C while crosses were kept at 29 °C. To prepare for crosses, 5-10 virgin *Drosophila* (Gal4 or UAS line) and 3-5 male *Drosophila* of the corresponding parent line were placed into plastic vials containing solid food. In general, Oregon-R was crossed with the specific GAL4 line of the particular analysis to produce GAL4-OreR whereas UAS-Aβ42 was crossed with the specific GAL4 line to produce the transgenic *Drosophila* line GAL4-Aβ42 that expressed Aβ42. Table 1 depicts the progeny lines used in different analyses.

**Table 1:**
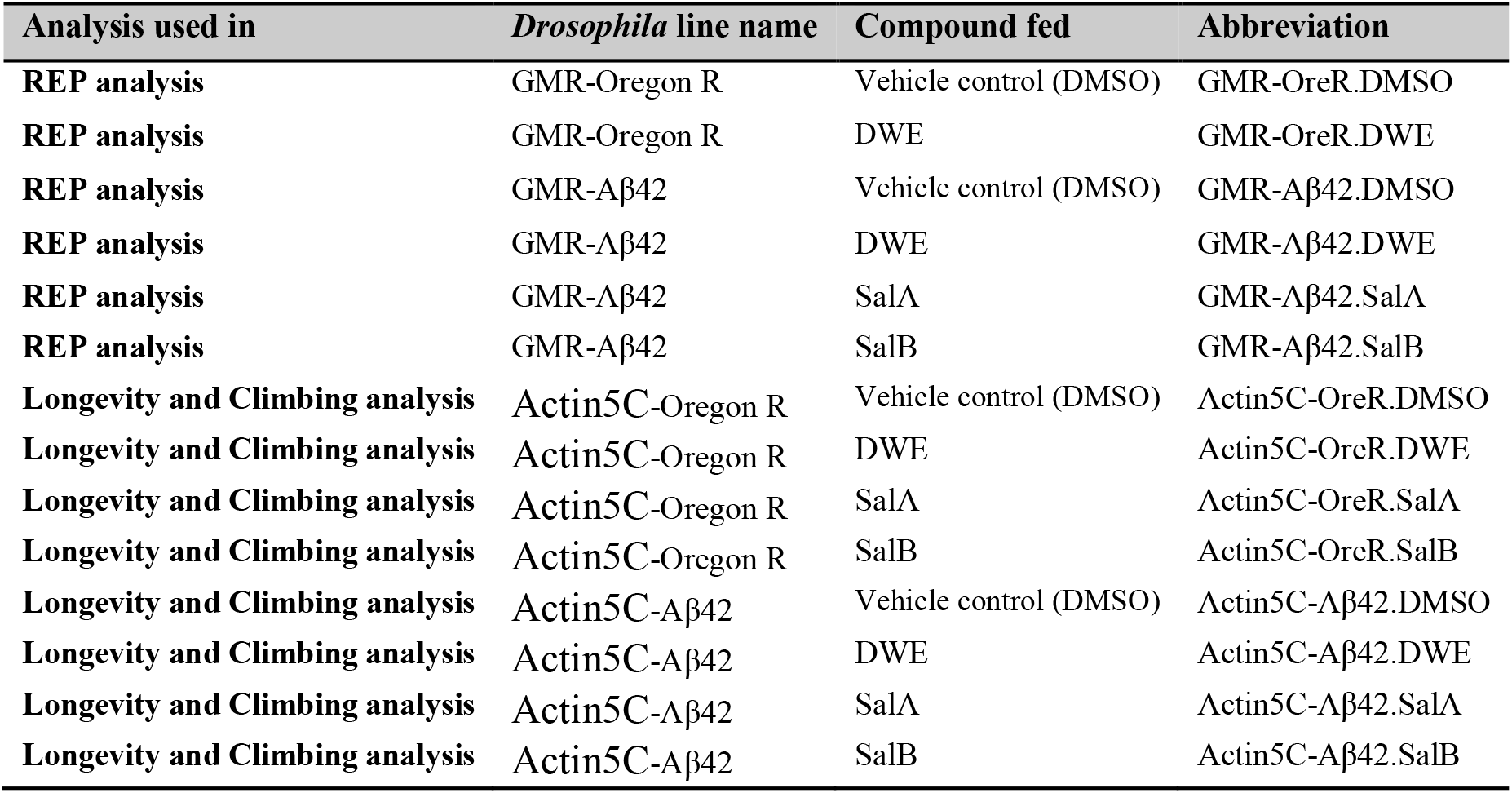
List of *Drosophila* lines used in corresponding studies

| Analysis used in | *Drosophila* line name | Compound fed | Abbreviation |
| --- | --- | --- | --- |
| REP analysis | GMR-Oregon R | Vehicle control (DMSO) | GMR-OreR.DMSO |
| REP analysis | GMR-Oregon R | DWE | GMR-OreR.DWE |
| REP analysis | GMR-Aβ42 | Vehicle control (DMSO) | GMR-Aβ42.DMSO |
| REP analysis | GMR-Aβ42 | DWE | GMR-Aβ42.DWE |
| REP analysis | GMR-Aβ42 | SalA | GMR-Aβ42.SalA |
| REP analysis | GMR-Aβ42 | SalB | GMR-Aβ42.SalB |
| Longevity and Climbing analysis | Actin5C-Oregon R | Vehicle control (DMSO) | Actin5C-OreR.DMSO |
| Longevity and Climbing analysis | Actin5C-Oregon R | DWE | Actin5C-OreR.DWE |
| Longevity and Climbing analysis | Actin5C-Oregon R | SalA | Actin5C-OreR.SalA |
| Longevity and Climbing analysis | Actin5C-Oregon R | SalB | Actin5C-OreR.SalB |
| Longevity and Climbing analysis | Actin5C-Aβ42 | Vehicle control (DMSO) | Actin5C-Aβ42.DMSO |
| Longevity and Climbing analysis | Actin5C-Aβ42 | DWE | Actin5C-Aβ42.DWE |
| Longevity and Climbing analysis | Actin5C-Aβ42 | SalA | Actin5C-Aβ42.SalA |
| Longevity and Climbing analysis | Actin5C-Aβ42 | SalB | Actin5C-Aβ42.SalB |

Solid cornmeal feed was prepared by boiling 4% (w/v) corn starch, 5% (w/v) polenta, 10% (w/v) brown sugar, 0.7% (w/v) agar, 5% (w/v) inactivated yeast, 3% (w/v) nipagin, 0.7% (v/v) propionic acid and 0.5% (v/v) DMSO either alone (vehicle control) or containing the concentration of compound indicated in the text with constant mixing, before being aseptically transferred into plastic vials to be cooled and solidified. Liquid feed for the CApillary FEeder (CAFE) assay did not include cornmeal or agar: 5% (w/v) yeast extract, 5% (w/v) glucose, 1.7% (w/v) tryptone, 3% (w/v) nipagin, 0.7% (v/v) propionic acid and 0.5% (v/v) DMSO either alone (vehicle control) or containing the concentration of compound indicated in the text.

Conversely, an altered version of liquid feed with the addition of tryptone (Ja et al., 2007) for the CApillary FEeder (CAFE) assay was prepared without cornmeal and agar: 5% (w/v) yeast extract, 5% (w/v) glucose, 1.7% (w/v) tryptone, 3% (w/v) nipagin, 0.7% (v/v) propionic acid and 0.5% (v/v) DMSO either alone (vehicle control) unless mentioned otherwise. Each capillary tube was filled with 10μL of liquid feed.

### External eye surface digital imaging and phenotypic analysis

Light microscopy images were viewed using the stereo-motorized light microscope model Olympus SZX16 (Olympus Optical) attached with an Olympus DP72 camera (Olympus Optical). Images were taken with CellSens Dimesion version 1.5 (Olympus Optical). All light microscopy images have maximum magnifications of 11x.

For scanning electron microscopy (SEM), ten progenies from each group were fixed overnight at 4°C in McDowell-Trump fixative (Sigma-Aldrich), containing 4% formaldehyde and 1% glutaraldehyde in 0.1 M phosphate buffer (Sigma) (pH 7.2). The specimens were washed in phosphate buffer three times before being post-fixed in 1% (w/v) osmium tetroxide (Sigma-Aldrich) at 25°C for an hour. The specimens were then washed with distilled water and dehydrated in a series of ethanol; 50%, 75%, 95% and 100% ethanol for 15 minutes each. The dehydrated specimens were immersed in hexamethyldisilazane (HMDS) (Sigma) for 10 minutes. The specimens were air-dried in a desiccator overnight. Dried specimens were then mounted, and gold coated to be viewed with SEM (SU8010; Hitachi Ltd., Tokyo, Japan).

Degree of ommatidia distortion was obtained from each image using a computational method called *Flynotyper* (https://flynotyper.sourceforge.net) through Image J that calculates a phenotypic score (P-value) (Iyer et al., 2018, 2016).

### CAFE Assay

The CApillary FEeder (CAFE) assay (Ja et al., 2007) was modified and used throughout both the lifespan analysis and locomotive analysis. Using 15 mL falcon tubes, the bodies of the tubes were drilled with 1 mm diameter holes while four 2 mm diameter holes were drilled on the base of the caps to allow insertion of truncated 10 uL pipette tips. Glass capillary tubes (Vitrex, Germany, Cat no.: 161313) filled with liquid feed by capillary action was inserted through the cap via the pipette tips. Capillaries were replaced every day. To facilitate egg laying, 1 mL of agar was added to all tubes.

### Lifespan Analysis

For survival analysis, Actin5C-Aβ42 *Drosophila* were collected within 24 hours from eclosion and transferred to the CAFE assay at standard density (less than 20 per vial) at 29°C and 60% humidity. Dead *Drosophila* were counted daily. Surviving *Drosophila* were flipped to new vials and liquid feed with respective compounds was changed every day. The median lifespan corresponds to the day at which 50% of the *Drosophila* in a cohort is alive. The maximum lifespan is the day at which the last *Drosophila* in a cohort dies. To eliminate any variation caused by cytoplasmic background effects, all crosses were set up with female virgins from Actin5C-Gal4. Each line was done in triplicates (n≃50 in each replicate). Lifespans of each line were compared to Vehicle control (Actin5C-Aβ42.DMSO) line and tested for significance with log-rank test.

### Measurement of Active compounds

Ultra-Performance Liquid Chromatography (UPLC) analysis was executed using an Acquity Ultra Performance LC series (Waters) with an autosampler and an Acquity UPLC PDA UV detector (Waters) with an Acquity UPLC Column (Waters) (50 × 2.1 mm). Analytical conditions consist of mobile phase A of 0.5% formic acid dissolved in water while mobile phase B had an acetonitrile: water ratio of 95:5, at flow rate of 0.5 mL/ min and signals were detected at 288 nm. The injection volume for all experiments was 2 μL. The gradient mode was implemented as follows: 5-20% of mobile phase B from 0 to 10 min, 20-25% of mobile phase B from 10 to 17 min and 25-55% of mobile phase B from 17 to 30 min.

SalA and SalB concentrations were measured by UPLC. Briefly, a hundred adult *Drosophila* were decapitated and immersed in Ultra Pure H2O (Mayhems Solution Ltd). For food analysis, 2 mL of food was dissolved in Ultra Pure H2O (Mayhems Solution Ltd). For faecal analysis, a Q-tip was used to pick up faeces around the tube wall. The Q-tip was then immersed in Ultra Pure H2O (Mayhems Solution Ltd) for sample collection. All samples were homogenized for 5 minutes before freezing in −80 °C for 2 hours. Frozen samples were freeze dried for 2 to 3 days until all traces of H2O was gone. Samples were resuspended with suitable solvent and centrifuged to remove pellet. The supernatant was used for UPLC analysis. All standards were solubilized in Ultra Pure H2O (Mayhems Solution Ltd) at 1 mg/mL.

### Negative Geotaxis Assay

Mobility of experimental *Drosophila* was measured in average climbing speed. The *Drosophila* were then tapped down on to the bottom of the vial at least 3 times before being allowed to scale the vial walls for 10 seconds. The videos of the *Drosophila* climbing up the vials were recorded and were analysed using the software ToxTrac (https://sourceforge.net/projects/toxtrac/) (Rodriguez et al., 2017, 2018). Examples of the videos can be found in the supplementary data.

## Results

### DWE, SalA and SalB reduced aggregation of Aβ42 *in vitro*

To verify whether DWE and its components SalA and SalB influence the aggregation of Aβ42, they were first subjected to a primary screening via Thioflavin T (ThT). ThT binds to fibrillated aggregates whereby the dye experiences a characteristic red shift of its emission spectrum (Groenning, 2010). At specific time points, the fluorescence generated by ThT binding to amyloid fibrils were then measured and quantified. Decreasing fluorescence of ThT signified the test compound’s ability to inhibit Aβ42 aggregation. Morin was added as a positive control in the assay (Kapoor & Kakkar, 2012).

At the time point of 3600 seconds, Morin reduced the RFU readings by 71.2% compared to DMSO. At the same time point, addition of DWE to Aβ42 peptides decreased RFU readings by 36.9% while Aβ42 peptides incubated with SalA and SalB experienced 65.9% and 50.8% decrement respectively in RFU readings when compared to the DMSO (Figure 1). This showed that DWE, SalA and SalB affected the aggregation of Aβ42 by reducing the fibrillation.

**Figure 1:**
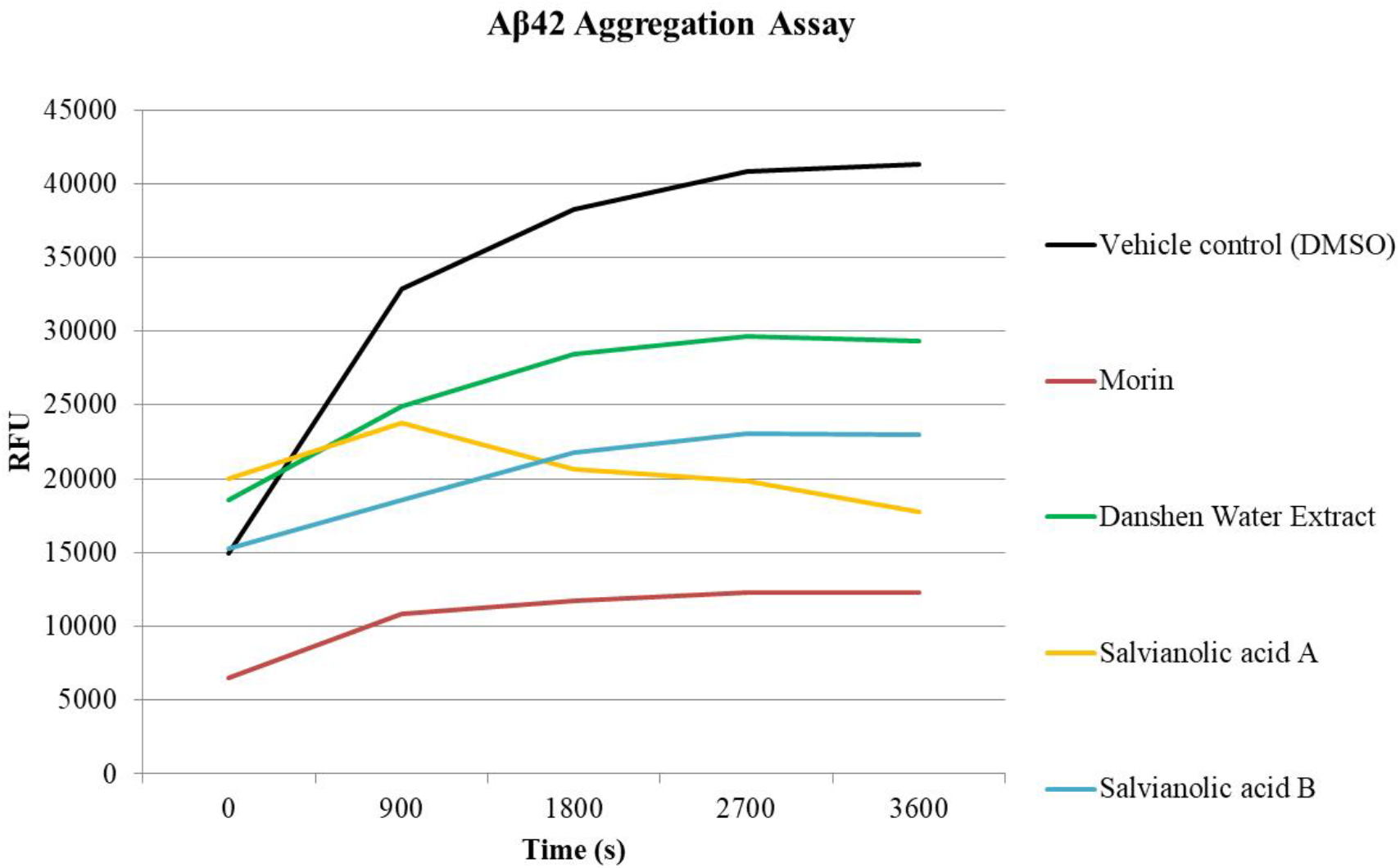
THT Aβ42 aggregation *in vitro* assay of DWE.

### DWE, SalA and SalB protected PC12 cells from Aβ42-induced cell death

To test the efficacy of DWE, SalA and SalB in cells, we tested the compounds using PC12 rat pheochromocytoma cells. The presence of Aβ42 peptides decreased PC12 cell viability to 40% compared to the unexposed control (p=0.0022). However, this toxic effect was rescued with the supplementation of DWE and its components, SalA and SalB (Figure 2). The best rescue effect was exhibited by incubation with SalB in which cell viability was increased to 95.4% cell viability based on the control when compared to the Aβ42-incubated PC12 cells supplemented with vehicle control (p=0.004). This was followed by SalB with a cell viability of 84% (p=0.0093) and DWE with cell viability of 73.5% (p=0.011). This demonstrated the ability of DWE, SalA and SalB to protect neuronal cells from the neurotoxicity effect of Aβ42.

**Figure 2:**
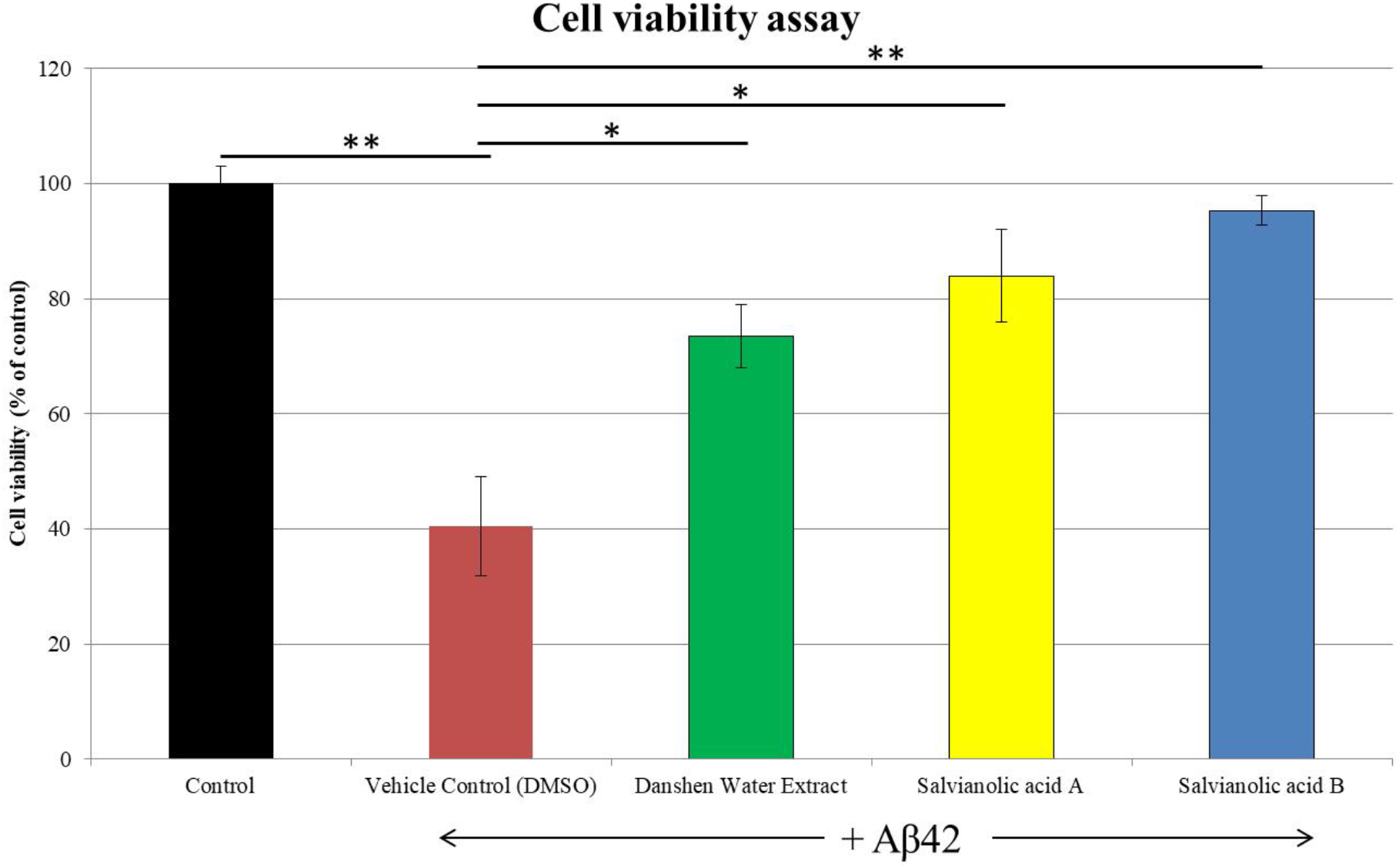
Cell viability assay of PC12 cells without addition of Aβ42 and compounds, and with addition of Aβ42 supplemented with vehicle control (0.5% DMSO), SalA or SalB. *P*-values indicated significance at *P<0.05, **P<0.005.

### SalA and SalB were detected in the brains and bodies of *Drosophila* after DWE feeding

In order to test the effect of these compounds on a whole organism, we chose *Drosophila melanogaster* as our model organism. As SalA and SalB are the two most abundant components in DWE (Ai & Li, 1988; Lian-niang et al., 1984), their presence in various parts of DWE-fed *Drosophila* were analysed using UPLC. The retention time of SalA and SalB were 21.9 minutes (Figure 3Ai) and 20.7 minutes (Figure 3Aii) respectively. By comparing the retention time of both reference standards, DWE with a concentration of 10mg/ml were found to have 43.0 μg/ml of SalA and 571 μg/ml of SalB.

**Figure 3:**
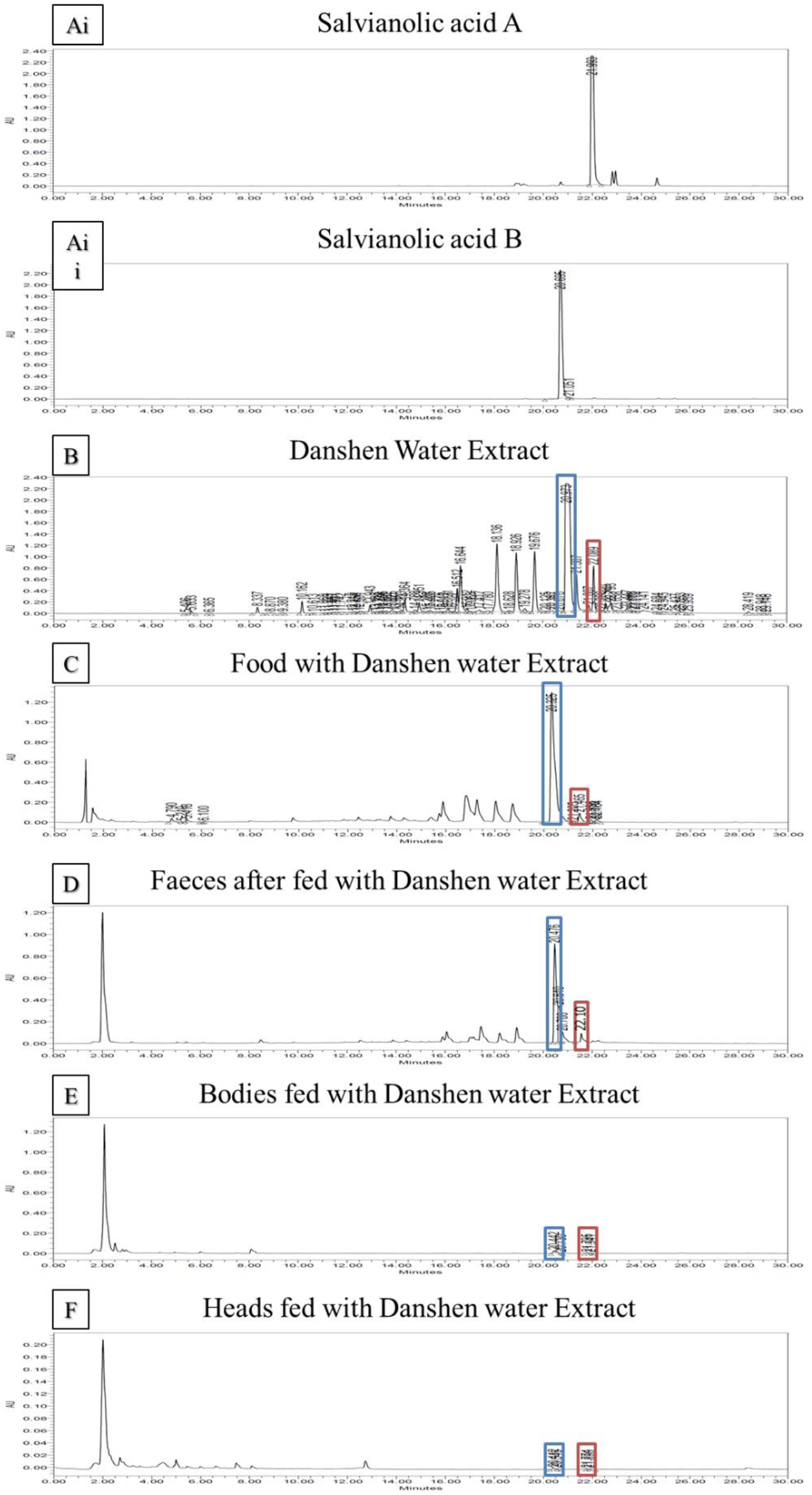
UPLC analysis of SalA and SalB. **Ai** and **Aii** depicts the standard graphs of pure SalA and SalB, respectively. **B.** shows the graph of DWE. **C.** shows the graph for food prepared with DWE while **D.** shows the graph of *Drosophila* faecal matter after fed food with DWE**. E.** and **F.** denote the graphs taken of the *Drosophila* bodies and heads, respectively after fed food with DWE. Blue boxes indicate SalB peaks while red boxes indicate SalA peaks.

Subsequently, one hundred *Drosophila* were fed with DWE and the heads, bodies and faeces were harvested and analysed through UPLC. For SalA, 0.85 μg/ml was detected in the heads, 4.3 μg/ml in the bodies and 37.3 μg/ml in the faeces. Likewise, the concentration of SalB in DWE-fed Drosophila heads, bodies and faeces were determined to be 2.6 μg/ml, 12.9 μg/ml and 407.1 μg/ml, respectively. The data confirmed the presence of SalA and SalB in the heads and bodies of *Drosophila* after feeding of DWE.

### Effect of DWE, SalA and SalB on *Drosophila melanogaster* AD model

The rough eye phenotype (REP) screening system was used to observe the effects of DWE, SalA and SalB on *Drosophila*, the rough eye phenotype (REP) (Kumar, 2012). The *Drosophila* eye is a model system to understand developmental neurobiology due to its simple neuroectoderm structure consisting of photoreceptors and accessory cells. Each eye comprised 800 hexagonal-shaped components known as ommatidium, organized in a crystalline array akin to the honeycomb cells of a beehive. These ommatidia are positioned in columns across the eye resulting in a concave “egg-like” formation (Figures 5A and 5A’). The mechano-sensory bristles extending at alternating vertices of each ommatidium that are directed at precise angles give an additional sensory field (Cagan, 2009; Cagan & Ready, 1989; Kumar, 2012). As each ommatidium contains seven photoreceptor nerve cells, any distortion in the eye morphology could be attributed to abnormalities in the neurons (Basler et al., 1991; Tomlinson et al., 1987).

By using light micrographs, a strongly observable REP was seen when human Aβ42 was ectopically expressed in the *Drosophila* eyes via the pan-retinal GMR-GAL4 driver (Figures 3B and 3B’) (Finelli et al., 2004). When compared to the control GMR-OreR.DMSO (Figures 4A and 4A’), the GMR-Aβ42.DMSO adult eyes were severely malformed with merged ommatidia (Figures 4B and 4B’)

**Figure 4:**
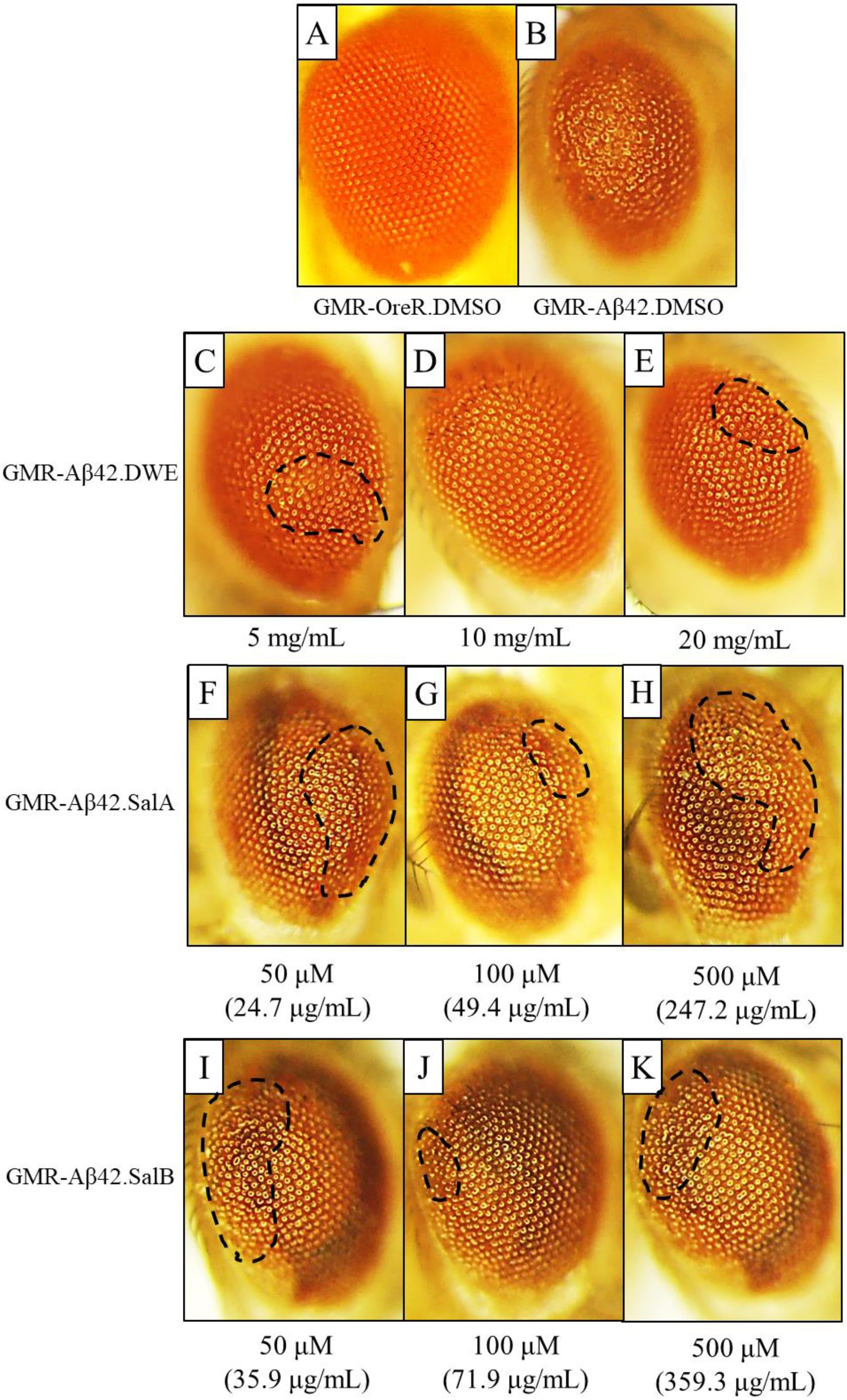
Light micrographs of transgenic *Drosophila melanogaster* eyes at magnifications x11.5. Black dotted circles indicate area with dense number of fused ommatidia. **A.** Eye of control GMR-OreR.DMSO. **B.** Eye of GMR-Aβ42.DMSO. **C.** to **E.** Eyes of GMR-Aβ42.DWE *Drosophila* fed food with 5 mg/mL, 10 mg/mL and 20 mg/mL of DWE, respectively. **F.** to **H.** Eyes of GMR-Aβ42.SalA *Drosophila* fed food with 50 μM, 100 μM and 500 μM of SalA, respectively. **I.** to **K.** Eyes of GMR-Aβ42.SalB *Drosophila* fed food with 50 μM, 100 μM and 500 μM of SalB, respectively.

The supplementation of transgenic *Drosophila* with different concentrations of DWE resulted in the partial rescue of eye deformation compared to that of GMR-Aβ42.DMSO albeit at varying degrees (Figures 4C-E). As the dosage of DWE increased, the rescue effect on the eyes of the Aβ42-expressing *Drosophila* fed with 10 mg/mL (Figure 4D) was more apparent showing similarity to those of the control GMR-OreR.DMSO (Figure 4A). However, the dosage of 20 mg/mL did not give an observable improvement on the eye (Figures 4E). Similarly for both SalA (Figure 4F-3H) and SalB (Figure 4I-3K) treatments, there was a recovery in eye morphology when transgenic Drosophila were cultured in 100 μM of compounds compared to 50 μM with 100 μM fed eyes having less fused ommatidia (indicated by dotted circles) and 500 μM feeding did not improve rectification of the eyes further. Thus, the optimum concentration for amelioration of the REP in GMR-Aβ42 transgenic flies for DWE, SalA and SalB are 10 mg/mL, 100 μM and 100 μM, respectively.

Scanning electron microscopy (SEM) was used to analyse the phenotype at higher magnification (Figure 5). GMR-Aβ42.DMSO eyes (Figure 5B and 5B’) showed merged and bulged ommatidia that had perforated holes which gave the eye a “glazed” exterior compared to GMR-OreR.DMSO’s (Figure 5A) “egg-like” shaped eye. In addition, the overall inter-ommatidial bristles were fewer than the Control and exhibited distorted polarities.

**Figure 5:**
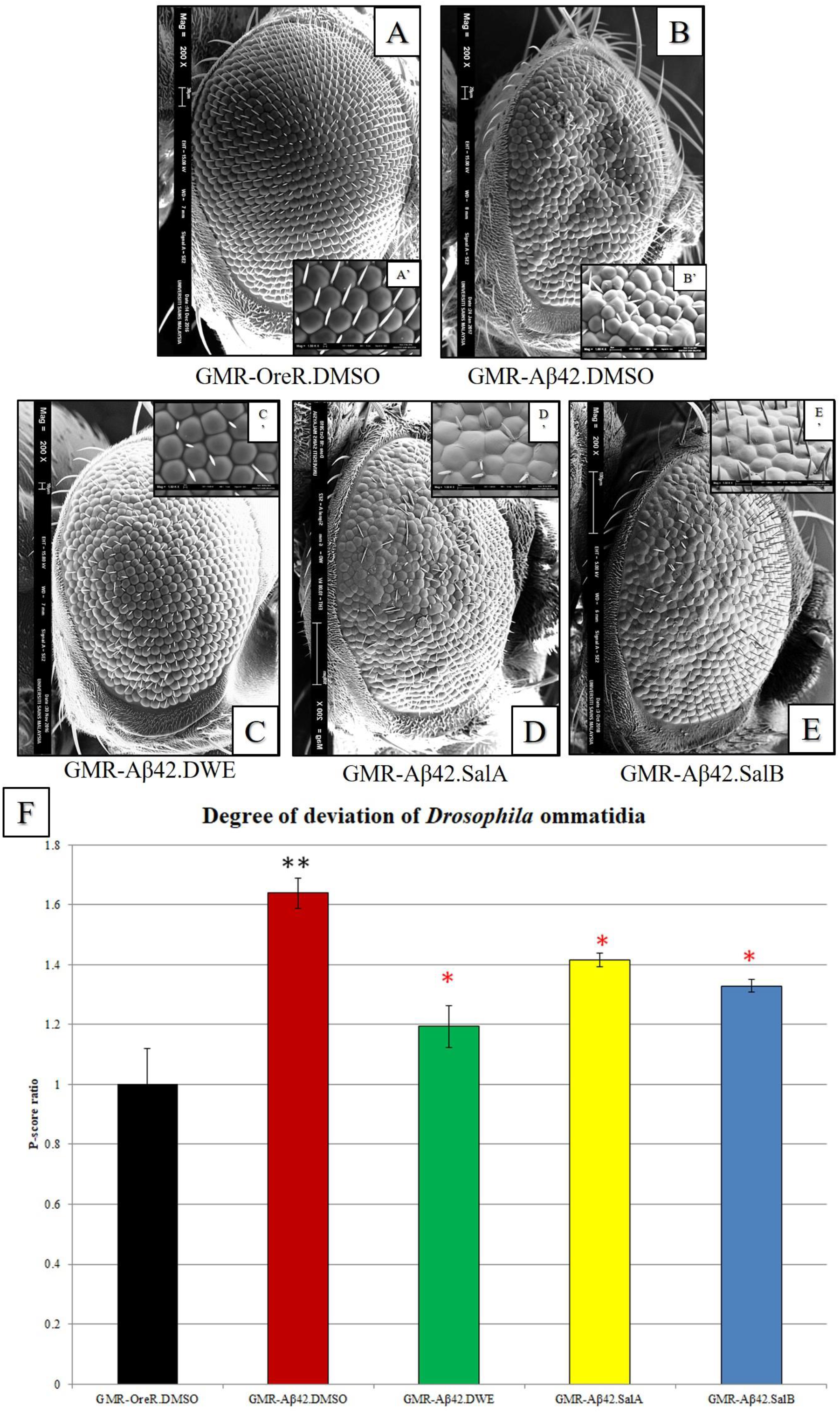
Scanning electron micrographs of transgenic *Drosophila melanogaster* at magnifications x200 and x1500. **A.** Eye of control GMR-OreR.DMSO. **B.** Eye of GMR-Aβ42.DMSO. **C.** Eye of GMR-Aβ42.DWE *Drosophila* fed food with 10 mg/mL of DWE. **D.** Eye of GMR-Aβ42.SalA *Drosophila* fed food with 100 μM SalA. **E.** Eye of GMR-Aβ42.SalB *Drosophila* fed food with 100 μM SalB. **F.** P-scores of the transgenic *Drosophila melanogaster* obtained from the *Flynotyper* software. *P*-values indicated significance at *P<0.05. Black asterisk (*) represents P-values against the control while red asterisk 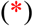 represents P-values against DMSO control

The extent of rescue effect on Aβ42’s toxicity was assessed using the *Flynotyper* software (Iyer et al., 2018, 2016). The quantification of morphological disfigurements in the *Drosophila* eye was evaluated in the form of the phenotypic score (P-score). The higher the P-score, the more distorted that specific eye is, and thus the more severe the REP is (Figure 5F). Parallel to SEM images (Figures 5A-G and 5A’-G’), the P-score for GMR-Aβ42.DMSO was significantly higher (p=0.0088) than GMR-OreR.DMSO, demonstrating Aβ42’s adverse effects on *Drosophila* eyes when not supplemented with any extra extracts or compounds. Feeding of DWE, SalA and SalB reduced the P-score on the eyes with GMR-Aβ42.DWE having the lowest P-score among compound-fed *Drosophila* compared to GMR-Aβ42.DMSO (p=0.0028), followed by SalB (p=0.0015) and SalA (p=0.011). These results implied that DWE and its components SalA and SalB showed amelioration towards Aβ42-induced neurodegeneration in *Drosophila* in a dosage dependent manner, with 10 mg/mL DWE being the optimum dosage.

### DWE, SalA and SalB extended the lifespan of the *Drosophila* AD model

The effect of prolonged exposure to 10 mg/mL DWE, 100 μM SalA and 100 μM SalB and their influence on longevity were investigated. In order to do this, the Actin5C-GAL4 driver which drives ubiquitous expression in muscle tissue was employed. There was no significant difference when Actin5C-OreR *Drosophila* was fed with DWE, SalA and SalB versus the same line cultured with only vehicle control (0.5% DMSO) for both males (Table 2, Figure 6A) and females (Table 3, Figure 6C). This indicated that DWE and its compounds did not exert any ill effects on the lifespan of control *Drosophila*. On the other hand, Actin5C-Aβ42.DMSO *Drosophila* was significantly different from Actin5C-OreR.DMSO with average median lifespans of male and female dropping by 42.8% and 47.6% respectively. Moreover, Actin5C-Aβ42 *Drosophila* lines with DWE, SalA and SalB consumptions were significantly different compared to Actin5C-Aβ42.DMSO. The average median lifespans for male Aβ42-expressing *Drosophila* increased by 75%, 37.5% and 37.5% after feeding of 10 mg/mL DWE, 100 μM SalA and 100 μM SalB respectively when compared to vehicle control-fed Actin5C-Aβ42.DMSO (Table 2, Figure 6B). Likewise, Actin5C-Aβ42.DMSO females experienced 63.6%, 36.7% and 72.7% extension in average median lifespan after being cultured in 10 mg/mL DWE, 100 μM SalA and 100 μM SalB respectively (Table 2, Figure 6D).

**Table 2:** Mean and median of male experimented *Drosophila melanogaster* lines fed with or without DWE, SalA or SalB

| Name | No. of subjects | Restricted mean |  |  | Age in days at % mortality |  |  |  |  |  |
| --- | --- | --- | --- | --- | --- | --- | --- | --- | --- | --- |
|  |  | Days | Std. error | 95% C.I. | 25% | 50% | 75% | 90% | 100% | 95% Median C.I. |
| Actin5C-OreR.DMSO | 156 | 17.14 | 0.76 | 15.65 ~ 18.63 | 10 | 14 | 24 | 31 | 41 | 13.0 ~ 15.0 |
| Actin5C-OreR.DWE | 182 | 17.77 | 0.73 | 16.35 ~ 19.19 | 10 | 15 | 26 | 33 | 41 | 13.0 ~ 16.0 |
| Actin5C-OreR.SalA | 156 | 17.14 | 0.76 | 15.65 ~ 18.63 | 10 | 14 | 24 | 31 | 41 | 13.0 ~ 15.0 |
| Actin5C-OreR.SalB | 154 | 16.26 | 0.79 | 14.72 ~ 17.80 | 8 | 13 | 23 | 32 | 42 | 12.0 ~ 15.0 |
| Actin5C-Aβ42.DMSO | 161 | 8.81 | 0.31 | 8.20 ~ 9.42 | 6 | 8 | 11 | 15 | 19 | 8.0 ~ 8.0 |
| Actin5C-Aβ42.DWE | 168 | 15.50 | 0.60 | 14.32 ~ 16.68 | 9 | 14 | 21 | 28 | 33 | 13.0 ~ 15.0 |
| Actin5C-Aβ42.SalA | 163 | 12.21 | 0.52 | 11.20 ~ 13.22 | 7 | 11 | 16 | 22 | 35 | 10.0 ~ 11.0 |
| Actin5C-Aβ42.SalB | 199 | 13.33 | 0.55 | 12.26 ~ 14.40 | 7 | 11 | 19 | 25 | 38 | 10.0 ~ 12.0 |

**Table 3:** Mean and median of female experimented *Drosophila melanogaster* lines fed with or without DWE, SalA or SalB

| Name | No. of subjects | Restricted mean |  |  | Age in days at % mortality |  |  |  |  |  |
| --- | --- | --- | --- | --- | --- | --- | --- | --- | --- | --- |
|  |  | Days | Std. error | 95% C.I. | 25% | 50% | 75% | 90% | 100% | 95% Median C.I. |
| Actin5C-OreR.DMSO | 194 | 21.01 | 0.68 | 19.66 ~ 22.35 | 13 | 21 | 29 | 34 | 41 | 18.0 ~ 22.0 |
| Actin5C-OreR.DWE | 198 | 21.11 | 0.75 | 19.65 ~ 22.57 | 12 | 20 | 30 | 36 | 41 | 17.0 ~ 22.0 |
| Actin5C-OreR.SalA | 154 | 19.25 | 0.89 | 17.50 ~ 20.99 | 9 | 17 | 28 | 36 | 44 | 15.0 ~ 19.0 |
| Actin5C-OreR.SalB | 159 | 19.65 | 0.91 | 17.87 ~ 21.44 | 11 | 17 | 29 | 38 | 45 | 15.0 ~ 18.0 |
| Actin5C-Aβ42.DMSO | 153 | 11.48 | 0.39 | 10.71 ~ 12.24 | 8 | 11 | 15 | 19 | 23 | 10.0 ~ 11.0 |
| Actin5C-Aβ42.DWE | 171 | 18.78 | 0.62 | 17.56 ~ 20.00 | 12 | 18 | 25 | 30 | 36 | 16.0 ~ 19.0 |
| Actin5C-Aβ42.SalA | 159 | 16.10 | 0.65 | 14.83 ~ 17.37 | 10 | 15 | 22 | 28 | 34 | 14.0 ~ 16.0 |
| Actin5C-Aβ42.SalB | 154 | 19.87 | 0.74 | 18.42 ~ 21.32 | 13 | 19 | 26 | 32 | 44 | 18.0 ~ 20.0 |

**Figure 6:**
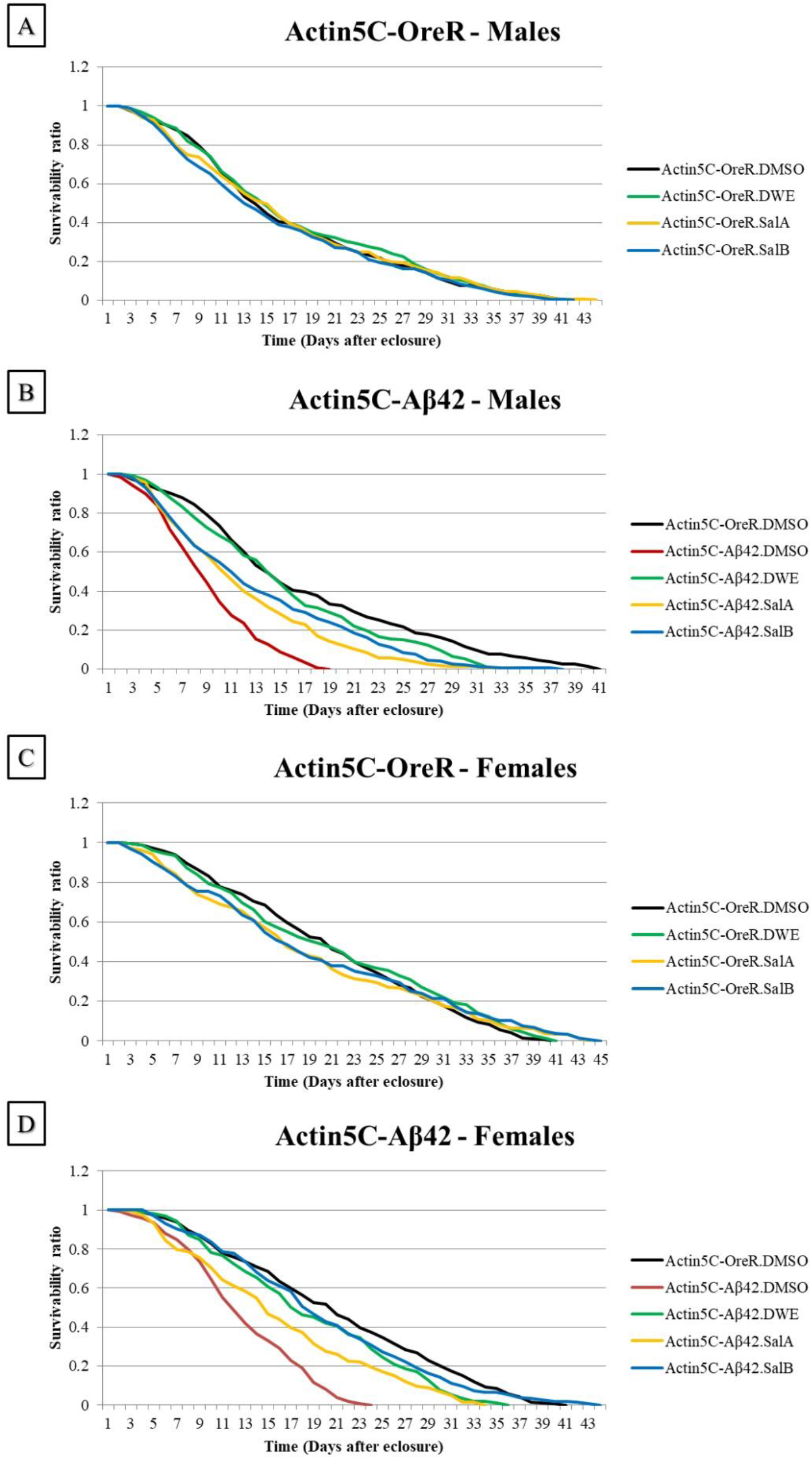
The effect of DWE on the lifespan of transgenic *Drosophila melanogaster*. **A.** and **B.** show the longevity graphs of Actin5C-OreR males and females, respectively. **C.** and **D.** depict the graphs of Actin5C-Aβ42 males and females, respectively.

To investigate the magnitude of rescue effect of the compounds, comparison of the ratio of Actin5C-Aβ42.compound:Actin5C-Aβ42.DMSO was made. As the same Actin5C-Aβ42.DMSO triplicate line was utilized to compare all of the compound-fed fly lines, the length of lifespan for Actin5C-Aβ42.DMSO remained static. The determining element depended on the lifespan length of compound-fed lines. Longer lifespans of compound-fed fly lines resulted in higher ratio values which indicated the compounds had a higher potency in prolonging lifespan. Based on the males, the ratios of Actin5C-Aβ42.DWE:Actin5C-Aβ42.DMSO for the restricted mean and average maximum lifespan were 1.75 and 1.74, respectively. For SalA, the ratios were 1.39 and 1.84, respectively while SalB males had ratios of 1.51 and 2.00, respectively. On the other hand, DWE females exhibited ratios of 1.64 and 1.57, respectively. Females of SalA had ratios of 1.40 and 1.48, respectively while SalB females showed ratios of 1.73 and 1.90. Here, we showed that DWE, SalA and SalB were able to prolong Actin5C-Aβ42 *Drosophila*’s severely shortened lifespan. When compared within sexes, DWE was the most effective in lengthening the lifespan of male Actin5C-Aβ42 *Drosophila*, followed by SalB and SalA. Alternatively, female Actin5C-Aβ42 *Drosophila* benefit most from SalB consumption followed by DWE and finally SalA.

### Increased average climbing speed (mm/s) in compound-fed AD *Drosophila*

In *Drosophila*, accumulation of Aβ42 peptides contributes to locomotor dysfunction (Iijima et al., 2004). Hence, the possibility of these compounds having significant effects on the mobility of the AD *Drosophila* was investigated. In this experiment, Actin5C-Aβ42 *Drosophila* were cultured with and without compounds (DWE, SalA and SalB) and the negative geotaxis assay was performed. This assay recorded the average climbing speed (mm/s) of the flies.

For male AD *Drosophila* fed with and without DWE (Figure 7A), there was a significant difference in the average climbing speed between Actin5C-Aβ42.DWE and Actin5C-Aβ42.DMSO. Actin5C-Aβ42.DWE had a higher average climbing speed than Actin5C-Aβ42.DMSO at all three time points. Similar to the males, there was a significant difference in the average climbing speed of female Actin5C-Aβ42.DWE and Actin5C-Aβ42.DMSO (Figure 7B). Actin5C-Aβ42.DWE showed higher average climbing speed than Actin5C-Aβ42.DMSO at all three time points.

**Figure 7.**
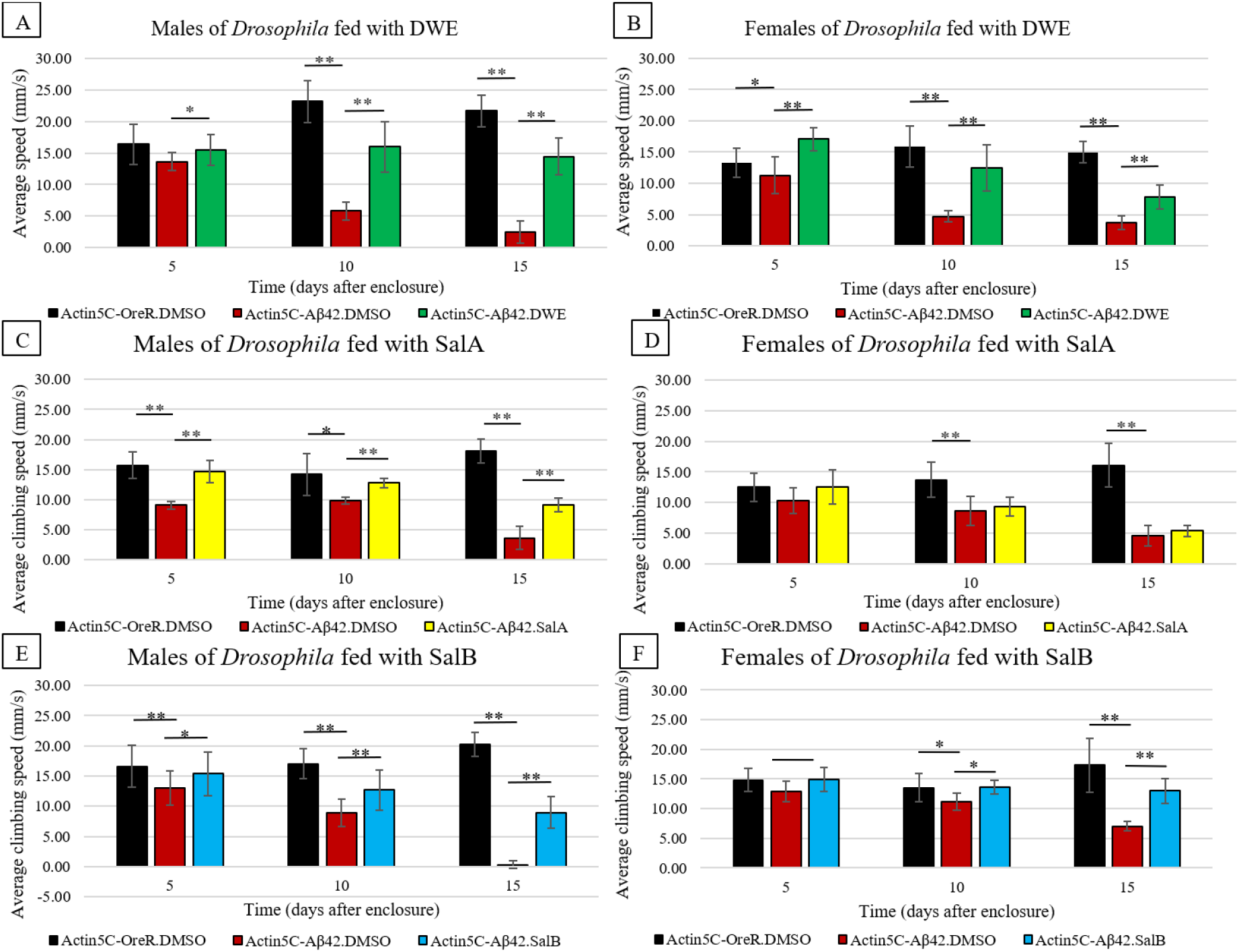
Average climbing speed (mm/s) of AD *Drosophila* fed with and without compounds at three time points (5 days, 10 days and 15 days). **A.** and **B.** show the average climbing speed of AD *Drosophila* fed with and without DWE. **C.** and **D.** show the average climbing speed of AD *Drosophila* fed with and without SalA. **E.** and **F.** show the average climbing speed of AD *Drosophila* fed with and without SalB. *P*-values indicated significance at *P<0.05 and **P<0.01.

For male AD *Drosophila* fed with and without SalA (Figure 7C), the male Actin5C-Aβ42.SalA showed significantly higher average climbing speed compared to Actin5C-Aβ42.DMSO at all three timepoints. Contrary to the male Actin5C-Aβ42.SalA (Figure 7D), female Actin5C-Aβ42.SalA showed no significant difference in average climbing speed between Actin5C-Aβ42.SalA and Actin5C-Aβ42.DMSO for all three time points.

For male AD *Drosophila* fed with and without SalB (Figure 7E), there was a significant difference between the Actin5C-Aβ42.SalB and Actin5C-Aβ42.DMSO for all three time points. Female Actin5C-Aβ42.SalB (Figure 7F) was also shown to have significant difference in average climbing speed when compared to Actin5C-Aβ42.DMSO but only on the 10^th^ and 15^th^ day.

To study the extent of rescue effect of the compounds, we compared the ratio between the average speed of AD *Drosophila* fed with and without the compounds, whereby a higher ratio indicates a better effect on the mobility. It was suggested that SalB had the best effect for male AD *Drosophila*. For the female counterpart, DWE had the best effect on all three time points, suggesting it to be the best compound even at an early stage.

## Discussion

With respect to the AD amyloidogenesis pathogenesis, neurotoxic amyloid plaques found in the brains of AD patients comprised primarily of a 40–42 amyloid-beta (Aβ) amino acid with Aβ42 being the most fibrillary species (Götz et al., 2011). The Chinese sage, Danshen has been employed throughout traditional Chinese medicinal history as a therapeutic agent for various cardiovascular and cerebrovascular diseases. This study takes advantage of Danshen’s vast beneficial properties aiming to uncover a potential remedy for Alzheimer’s disease. Here, Danshen and its water-soluble components SalA and SalB on Aβ42 were tested both in *in vitro* and in a model organism.

We first verified that DWE, SalA and SalB possessed the ability to reduce fibrillation rate of Aβ42 using the Aβ42 aggregation assay. SalA was found to be the most proficient inhibitor followed by SalB and lastly DWE. Indeed, SalA’s addition to Aβ42 peptides not only yielded lowered RFU readings but also led to decreasing RFU intensities from the 900th second. This could be due to its potential ability to revert the Aβ42 fibrils to their monomeric form. This was supported by previous work that exhibited the propensity of SalA to disaggregate pre-formed Aβ42 fibrils (Cao et al., 2013).

Next, the ability of these inhibitors to exert neuroprotective effect on cells in a controlled-setting was investigated. The pheochromocytoma PC12 cells obtained from rat adrenal medulla were chosen for their extremely high sensitivity to Aβ-associated neurodegeneration, a response higher than the human cell line, SH-SY5Y (Sakagami et al., 2017, 2018). When tested on the PC12 cells, SalB was most efficient in protecting PC12 cells against Aβ42’s neurotoxicity, followed by SalA and DWE. A key basis for the formation and stability of Aβ fibrils is the fitting and stacking of aromatic rings between the peptides that provide strong π –π interactions, thus stabilizing the complex (Chini et al., 2009; Makin et al., 2005). DWE contains a high amount of polyphenols with SalA and SalB being the most abundant. Polyphenols have been shown to weaken the aromatic links between Aβ fibrils (Porat et al., 2006). However, DWE was the least effective inhibitor in this assay which could be attributed to it being a crude extract consisting of various components that might block the binding of polyphenols available in the extract when tested in an *in vitro* environment. In contrast, compounds SalA and SalB were over 98% pure.

Subsequent experiments were carried out with live organisms. Prior to behavioural studies, investigations on whether SalA and SalB were digested by the *Drosophila* as well as determination of the parts of the body that the compounds were transported to were performed. It was observed that while most of both compounds were expelled out of the *Drosophila*, there were small amounts detected in the bodies and heads. This showed the possibility that there was a transfer of both compounds to the brain after the feeding of DWE. It also raised the likelihood that SalA and SalB were utilized by neurons in live organisms. However, there were undetectable levels of SalA and SalB in the heads, bodies and faeces which could be due to their short half-life (Pei et al., 2008, Y. T. Wu et al., 2006).

The neurotoxicity assay that uses the eye structures of the *Drosophila* showed that DWE was the most effective treatment against Aβ42-induced REP followed by SalB and SalA. The longevity assay exhibited similar results with slight variations between genders. SalB worked best for females followed by DWE and SalA. In *in vitro* condition, SalA was the best inhibitor of Aβ42 aggregation. However, it provided the least protection when fed to a whole organism and this could be due to its incomplete cellular absorption compared to SalB. This difference in absorption was previously demonstrated in rats whereby oral administration of SalA showed a significantly lower plasma concentration of only 308 ng/ml compared to SalB which reached 1.5 μg/ml (Y. T. Wu et al., 2006, Pei et al., 2008). The UPLC findings from this study further supported this claim with the concentration of SalB found in the brains of DWE-fed flies being three times more than that of SalA.

The negative geotaxis assay to assess mobility recovery suggested that the compounds have a rescue effect on most of the AD *Drosophila* with the exception of females fed with SalA. While the exact mechanism is still unknown, this protective effect could be due to the antioxidative properties of the compounds as PC12 cells were extremely receptive to changes in oxygen levels (Alvarez-Tejado et al., 2001). It was shown by Li et al., (2008) that there was an elevated amount of reactive oxygen species/reactive nitrogen species (ROS/RNS) which preceded mitochondrial dysfunction, apoptosis and cell death when PC12 cells were exposed to Aβ (Guo et al., 2013). Mitochondrial damage prompted the loss of ATP (Li et al., 2008; Moreira et al., 2010, Chong et al., 2019) and resulted in a surge in ROS, further causing apoptotic cell death (Chong et al., 2019). In previous studies, salvianolic acids including SalA and SalB have been shown to inhibit the production of ROS in experimental stroke (Lv et al., 2015) and liver injury (Z. Wu et al., 2007) in rats. As such, this antioxidative mechanism of DWE and its polyphenol components might be functioning similarly for Aβ-incubated PC12 cells and AD *Drosophila*.

## Conclusion

DWE as a whole was able to rescue AD phenotypes in a sex-dependent manner when introduced to AD Drosophila. This is the first study at the time of writing that employed Drosophila melanogaster to study the neuroprotective effects of DWE and its components SalA and SalB on neuro-diseases, specifically AD. It is hoped that these discoveries will generate further queries into the fundamental aspects of the protective abilities of the extract and its compounds and thereby assist in the prevention of neuro-diseases in human.

## Acknowledgements

We would like to thank all our collaborators and colleagues for the discussion and the work conducted in this lab. We would also like to extend our gratitude to Emiko Sanada, Tatsuro Kawamura and Kaori Honda of RIKEN CSRS for their assistance in the project. This study was funded by the USM Top Down Research Fund – URICAS (1001/PBIOLOGI/870029). Florence Hui Ping Tan is the recipient of the MyBrainSc scholarship and RIKEN International Program Associate.

## Supplementary Data

The videos for the negative geotaxis assay can be downloaded from the link. The flies in each tube are in the order of (left to right): control (Actin5C-OreR.DMSO), AD *Drosophila* fed without compounds (Actin5C-Aβ42.DMSO) and AD *Drosophila* fed with compounds (Actin5C-Aβ42.DWE / Actin5C-Aβ42.SalA / Actin5C-Aβ42.SalB). (https://drive.google.com/open?id=181bmDjS935dIHDtvSmIB7jl3pjRcnWs2)

## Graphical Abstract

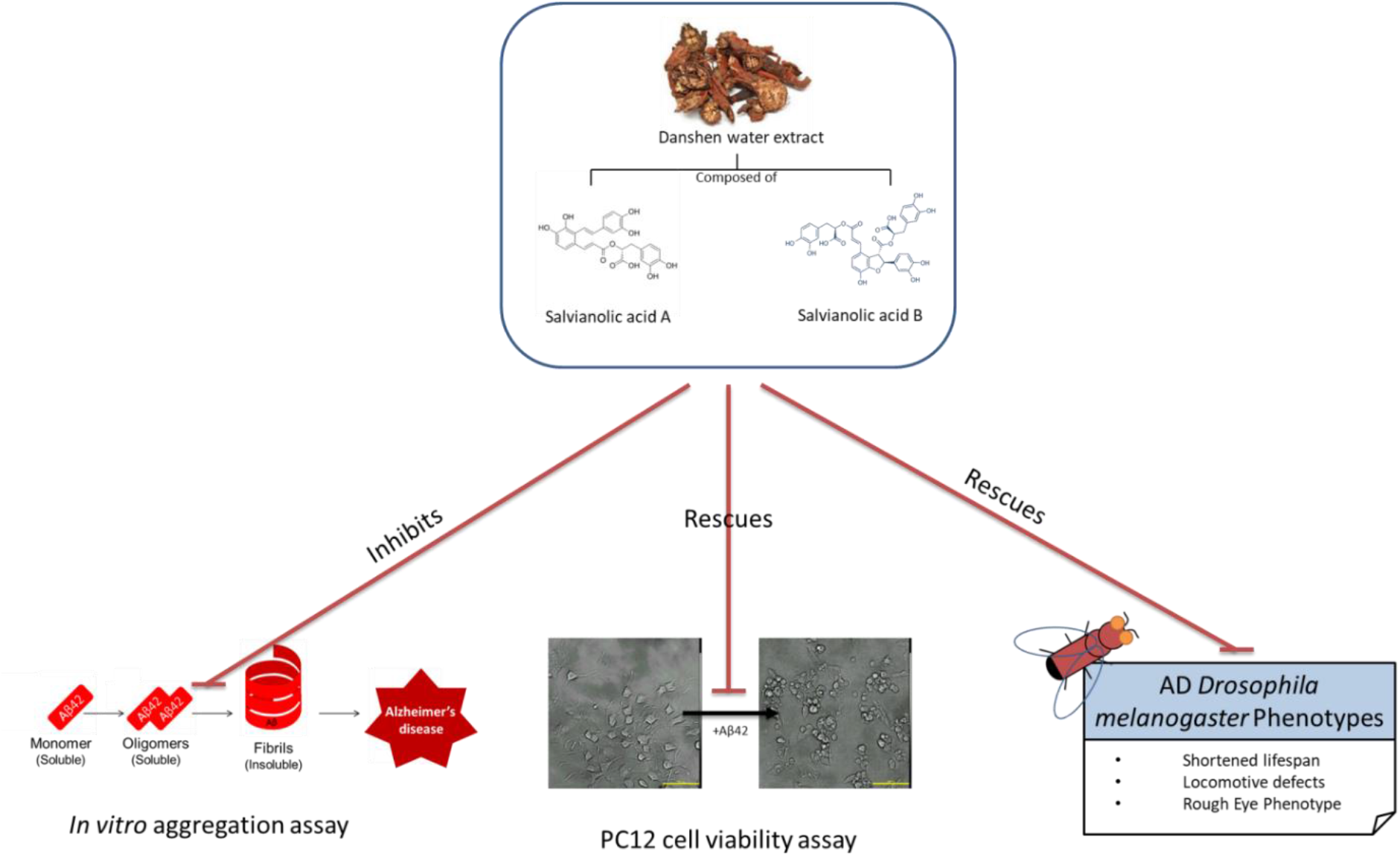

